# Resuscitation of the microbial seed bank alters plant-soil interactions

**DOI:** 10.1101/2020.06.02.130443

**Authors:** Venus Kuo, Brent K. Lehmkuhl, Jay T. Lennon

## Abstract

While microorganisms are recognized for driving belowground processes that influence the productivity and fitness of plant populations, the vast majority of bacteria and fungi in soil belong to a seed bank consisting of dormant individuals. Still, plant performance may be affected by microbial dormancy through its effects on the activity, abundance, and diversity of soil microorganisms. To test how microbial seed banks influence plant-soil interactions, we purified recombinant resuscitation promoting factor (Rpf), a bacterial protein that terminates dormancy. Then, in a factorially designed experiment, we applied the Rpf to soil containing field mustard (*Brassica rapa*), an agronomically important plant species. Plant biomass was ~33 % lower in the Rpf treatment compared to plants grown with an unmanipulated microbial seed bank. In addition, Rpf reduced soil respiration, decreased bacterial abundance, and increased fungal abundance. These effects of Rpf on plant performance were accompanied by shifts in bacterial community composition, which may have diluted mutualists or resuscitated pathogens. Our findings suggest that changes in microbial seed banks may influence the magnitude and direction of plant-soil feedbacks in ways that affect above- and below-ground biodiversity and function.

## INTRODUCTION

Belowground soil microbial communities play a critical role in determining the productivity and fitness of aboveground plants. Plant roots are intimately associated with thousands of bacterial and fungal taxa that are involved in soil processes such as decomposition, nutrient cycling, and pathogen suppression (Bardgett & van der Putten, 2014; Berendsen, Pieterse, & Bakker, 2012; Wagg, Bender, Widmer, & van der Heijden, 2014). In addition to containing mutualistic symbionts, soils harbor pathogenic microorganisms that reduce plant performance (Mansfield et al., 2012). Together, belowground microbial communities are responsible for generating plant-soil feedbacks that can influence the diversity and composition of plant communities (Bever, 2003). However, the direction and magnitude of soil microbial effects on plants is variable (Hoeksema et al., 2010; Kulmatiski, Beard, Stevens, & Cobbold, 2008) and likely influenced by the complexity of belowground communities (van der Heijden, de Bruin, Luckerhoff, van Logtestijn, & Schlaeppi, 2016; Wagg et al., 2014).

One belowground feature that is commonly overlooked when considering plant-microbe interactions is the metabolic heterogeneity of soil microbial communities. Many microorganisms are capable of entering a reversible state of reduced metabolic activity (i.e., dormancy) when challenged by suboptimal environmental conditions (Lennon & Jones, 2011). A large proportion (≥90 %) of microorganisms are dormant in soils (Alvarez, Alvarez, Grigera, & Lavado, 1998; Blagodatskaya & Kuzyakov, 2013; Blagodatsky, Heinemeyer, & Richter, 2000; Lennon & Jones, 2011). Collectively, these inactive individuals create a seed bank, which not only maintains biodiversity, but also affects ecosystem functioning. The incorporation of microbial dormancy into ecosystem experiments and models has been shown to improve predictions for soil microbial activity and nutrient cycling (Salazar, Lennon, & Dukes, 2019; Wang et al., 2015). Yet, the influence of microbial seed banks on plant-soil interactions remains to be determined.

Microbial seed banks may prevent local extinctions when exposed to fluctuating and stressful conditions that are typical in soil habitats (Shoemaker & Lennon, 2018). If microbial mutualists can persist by entering a seed bank, then this may promote beneficial soil functions such as disease-suppression, induction of plant immune response, stimulation of root growth, and biofertilization, which together, can increase plant performance (Lugtenberg & Kamilova, 2009; Mendes et al., 2011; Peralta, Sun, McDaniel, & Lennon, 2018; Wagg et al., 2014). In contrast, seed banks may contain pathogenic microorganisms that can reduce plant performance. For example, it is well known that destructive pathogens can reside in soil for months to years after the demise of plants, rendering potentially arable soil unusable (Koike, Subbarao, Davis, & Turini, 2003; Peeters, Guidot, Vailleau, & Valls, 2013). Although microbial seed banks likely contain both mutualistic and pathogenic taxa, the net effect of dormancy transitions on plant performance is unknown.

Resuscitation is an essential process that regulates microbial seed-bank dynamics (Lennon, den Hollander, M., & Blath, 2020). Dormancy is often terminated via the interpretation of environmental cues or communication signals that are produced by other microorganisms (Dworkin & Shah, 2010). One such signal is a bacterial protein called resuscitation promoting factor (Rpf), a muralytic enzyme that degrades the β-(1,4) glycosidic bond in peptidoglycan, which is a major cell-wall component of virtually all bacteria (Mukamolova, Kaprelyants, Young, Young, & Kell, 1998). Rpf genes are broadly distributed among the G+C-rich Gram-positive Actinobacteria (Ravagnani, Finan, & Young, 2005; Schroeckh & Martin, 2006). In some soil samples, Rpf homologs can be found in up to 25 % of all genomes (Lennon & Jones, 2011). Rpf has cross-species effects that stimulate the growth of closely related dormant bacteria (Puspita, Kitagawa, Kamagata, Tanaka, & Nakatsu, 2015; Schroeckh & Martin, 2006) either through its direct effects on cell wall integrity or through the release of small peptide fragments that cross-link peptidoglycan, which may serve as signaling molecules (Dworkin, 2014). Under laboratory conditions, Rpf can stimulate the activity of some microbial taxa at picomolar concentrations, but it has also been shown to inhibit the growth of other microorganisms (Mukamolova et al., 1998; Mukamolova et al., 2002), which is not surprising given that Rpf is homologous to lysozyme (Cohen-Gonsaud et al., 2005), an antimicrobial enzyme produced by many eukaryotic organisms. Overall, Rpf may have complex and interactive effects that influence plant-soil interactions.

In this study, we explored how seed-bank dynamics affect plant-microbe interactions by manipulating the process of resuscitation. Throughout the lifespan of a host plant (*Brassica rapa*), we applied purified recombinant Rpf to soils containing a complex microbial community. The experiment was guided by a conceptual model that considers how active and inactive microorganisms interact and potentially create feedbacks with plants (Fig. 1). We hypothesized that resuscitating the microbial seed bank with Rpf would alter plant performance through changes in the abundance, activity, and composition of soil microbial communities. On the one hand, Rpf treatment could enhance plant performance by waking up mutualistic taxa in the soil microbial community. On the other hand, resuscitating dormant microorganisms could negatively influence plant performance by diluting the benefits of mutualistic taxa or through the recruitment of pathogenic microorganisms. Here, we evaluate these outcomes while considering other effects of Rpf on plants, microbes, and their interactions.

**Fig. 1.**
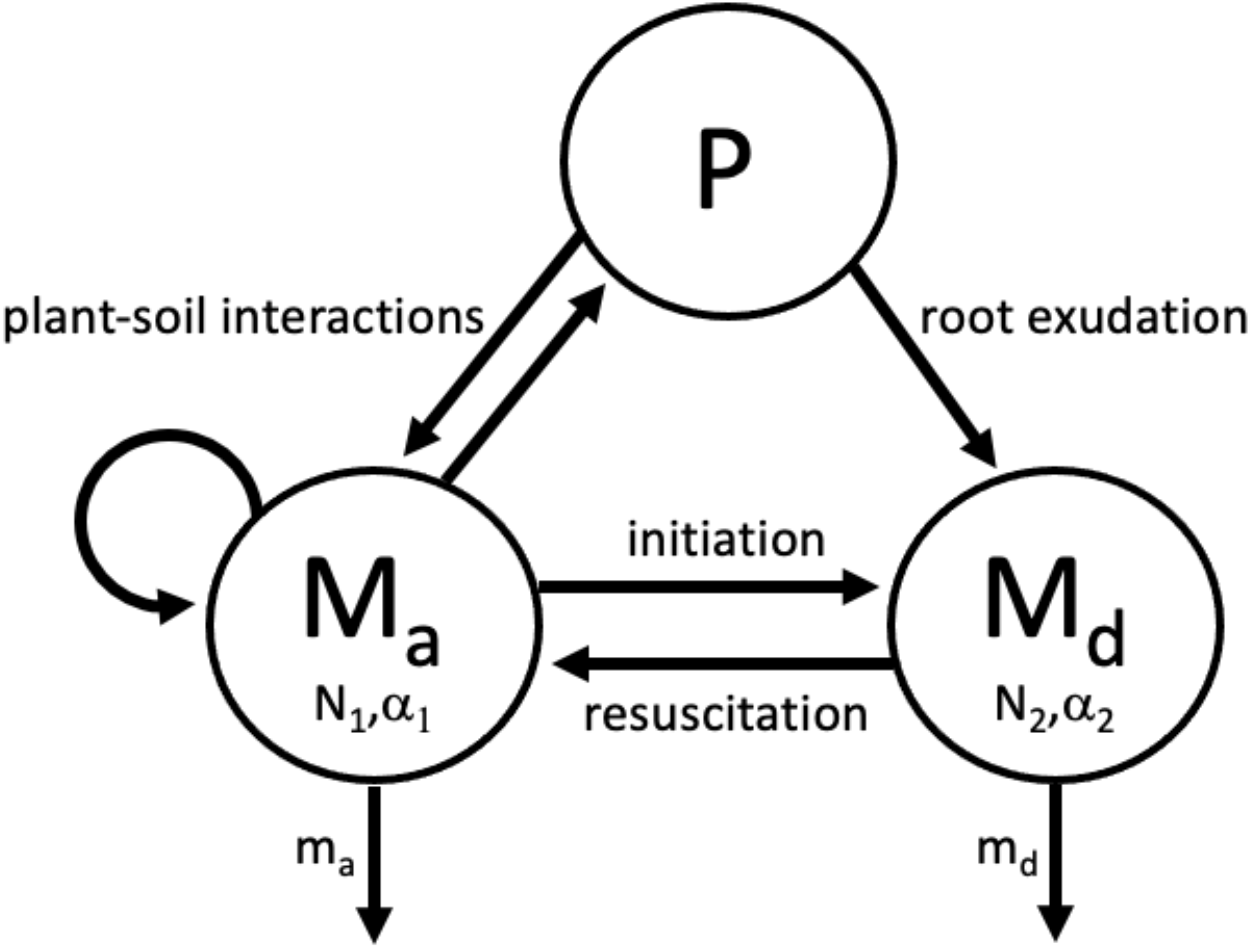
Conceptual framework depicting the role of the soil microbial seed bank in local plantmicrobe interactions. Members of the soil microbial community can transition between the metabolically active pool (M_a_) and the seed bank (M_d_) through initiation into dormancy and resuscitation into an active state. Both M_a_ and M_d_ pools have associated levels of microbial abundance (*N*) and diversity (*α*). While only members in M_a_ can reproduce, the baseline rate of mortality (*m_d_*) in M_d_ is reduced relative to the rate for M_a_ (*m_a_*). Active microorganisms (M_a_) and plants (P) can directly interact in the soil with one another through a suite of mechanisms, such as mutualism, parasitism, root colonization, as well as the secretion of phytochemicals or other signaling compounds. We envision that the M_d_ pool has minimal effects on plants. In contrast, this pool of inactive microorganisms may be resuscitated via processes including the release of root exudates, which may vary in space and time.

## METHODS

### Experimental design

We conducted a growth chamber experiment to test the effect of Rpf on plant-soil interactions using *Brassica rapa* L. (Brassicaceae), an economically important plant that consists of a variety of cultivated subspecies. We sowed 16 *B. rapa* seeds into individual pots, which were obtained from Wisconsin Fast Plants™ (Standard stock seeds, Wisconsin Fast Plants Program, University of Wisconsin, Madison, WI, USA). While *B. rapa* is non-mycorrhizal, it still associates with a diverse microbial community that influences plant growth (Lau & Lennon, 2012). For our experiment, we used plastic pots (17 cm diameter, 13 cm height) filled with an autoclaved (121 °C, 15 PSI, 16 h) substrate consisting of, on a volumetric basis, one part Metro Mix, one part Vermiculite, and one part soil. Based on a previous study, this mixture prevented compaction and promoted *B. rapa* growth (Lau & Lennon, 2011). The soil added to the potting mixture was collected from Indiana University Research and Teaching Preserve (DMS 39 °11’56.5”N 86 °30’40.5”W). We collected this silt-loam soil from the surface (0-5 cm) of a mesic, mid-successional site that was dominated by oaks (*Quercus velutina* and *Quercus rubra*) and sugar maple (*Acer saccharum*) within the Norman Upland bedrock physiographic unit (Schneider, 1966). Chemical and physical characteristics of the soil are listed in Table S1.

For the +Rpf treatment, we added 1 mL of recombinant Rpf protein (10 μM) to the surface of the soil substrate each week in three equidistant locations 5 cm away from main plant stem. The same was done for the-Rpf treatment (negative control), except we used protein buffer (20 mM Tris-HCl, 100 mM NaCl) instead of recombinant Rpf. Our experiment was conducted in a Percival model PGC-15 growth chamber with high efficiency light lamps (Philips series 700 32 watts 4100K F32T8/TL741 AltoII). The growth chambers were set to constant and full light intensity (1100 μmol m^-2^ s^-1^) with controlled humidity (60 %) and temperature (28 °C). Pots were arranged in a fully randomized design where each plant was assigned to a resuscitation treatment (i.e., -Rpf vs. +Rpf). All pots were watered with equal amounts of filtered and deionized water every other day.

### Recombinant Rpf

To manipulate microbial seed banks, we overexpressed Rpf from *Micrococcus luteus* sp. KBS0714, a bacterial strain isolated from agricultural soil at the W.K. Kellogg Biological Station, Hickory Corners, MI (MSU) (Kuo, Shoemaker, Muscarella, & Lennon, 2017; Lennon, Aanderud, Lehmkuhl, & Schoolmaster Jr, 2012). *Micrococcus luteus* sp. KBS0714 is a close relative of *M. luteus* NCTC 2662 (99 % sequence 16S rRNA sequence similarity, NCBI CP001628.1), a model organism used for studying Rpf (Mukamolova et al., 1998). To create a Rpf expression host, we amplified and cloned the *rpf* gene from KBS0714 into the expression vector pET15b. First, we extracted KBS0714 genomic DNA from pure culture using a Microbial DNA isolation kit (MoBio, Carlsbad, CA) for PCR amplification of the open reading frame of the *rpf* gene using two primers, Upper-F 5’ GCC CAT ATG GCC ACC GTG GAC ACC TG 3’ and Lower-R 5’ GGG GAT CCG GTC AGG CGT CTC AGG 3’, with incorporated restriction sites EcoRI, NdeI (forward primer) and BamHI (reverse primer) (Koltunov et al., 2010; Mukamolova et al., 1998). We amplified the *rpf* gene sequence with the following PCR conditions: initial: 95 °C for 5 min, 30 cycles of 95 °C for 30 s, 55 °C for 30 s, 72 °C for 1 min, and one final extension at 72 °C for 7 min. The *rpf* gene amplicon was ligated into pET15b (Invitrogen) as an EcoRI/BamHI fragment and transformed into *Escherichia coli* TOP10 (a non-expression host). We then identified positive clones with correct *rpf* sequence and orientation by Sanger sequencing at the Indiana University Center of Genomics and Bioinformatics (IU-CGB). Next, we extracted purified pET15b plasmids with the *rpf* gene from transformed TOP10 *E. coli* using the QIAprep Spin Miniprep Kit (Qiagen) following the manufacturer’s protocol. Finally, we transformed the purified pET15b expression vector with the *rpf* gene into the *E. coli* Origami BL21 (DE3) expression host incorporated with a polyhistidine-tag on the N-terminus of the recombinant protein.

To overexpress Rpf, we grew the *E. coli* expression host in Lysogeny Broth (LB) with appropriate antibiotics (ampicillin 100 μg mL^-1^, kanamycin 15 μg mL^-1^, and tetracycline 12.5 μg mL^-1^). During logarithmic growth, recombinant protein production was induced with Isopropyl β-D-1-thiogalactopyranoside (IPTG) (100 μM final concentration). We confirmed overexpression of Rpf with Western blots (Fig. S1). Cells were then collected by centrifugation, lysed by sonication, and filtered through a Ni-NTA Purification System (Invitrogen) using a 10 mL gravity fed column with a 2 mL resin bed to purify recombinant Rpf with the N-terminus polyhistidine-tag. Recombinant Rpf protein was washed with 5 mM imidazole buffer (300 mM NaCl, 50 mM Tris-HCl, 5 mM imidazole) and then eluted with 125 mM imidazole buffer. Rpf protein was purified by buffer exchange using a 10 mL Zeba Spin Desalting Columns (Thermo Fisher) with protein buffer (20 mM Tris-HCl, 100 mM NaCl) following manufacturer’s instructions and then passed through a 0.2 μm syringe filter before adding to soil substrate as described above.

### Plant responses

We censused the plants every day and recorded germination and flowering date. After most plants had finished flowering, we documented flower number, plant height, and specific leaf area (SLA = leaf area/dry mass) as previously described (Lau & Lennon, 2012). All open flowers in each experimental unit were hand-pollinated by other open flowers in the same treatment group with a soft paint brush, which was cleaned with 30 % isopropyl alcohol between treatment groups to prevent gene flow. Plants and seeds were harvested at the end of the six-week growing period when most individuals ceased flowering and had begun to senesce. As an estimate of female fitness, we removed plant seed pods and counted the number of seeds produced. To estimate male fitness, we counted the number of flowers per plant. In addition, we measured total, aboveground, and belowground biomass of each plant after drying plants at 65 °C for 48 h. We performed two-tailed Student’s *t*-test to evaluate the effect of Rpf on total biomass, aboveground (i.e., shoot) biomass, root biomass, shoot: root ratio, SLA, and plant fitness (i.e., flower and seed numbers).

In addition to our main study, we conducted a smaller scale experiment with *Arabidopsis thaliana*, a relative of *B. rapa* that also belongs to the Brassicaceae. Because this species is amenable to being grown axenically in the absence of soil, we were able to test for the direct effect of recombinant Rpf on plant performance. We placed seeds on sterile Murashige-Skoog (MS) agar plates containing either Rpf protein (final concentration: 1.6 μmol L^-1^) (+Rpf) or protein buffer control (-Rpf). For five weeks, we maintained eight seedlings in a Percival model PGC-15 growth chamber equipped with high efficiency lighting (Philips series 700 32 watts 4100K F32T8/TL741 AltoII). The growth chamber was set to constant and full light intensity (1100 μmol m^-2^ s^-1^) with controlled humidity (60 %) and temperature (28 °C). At the end of the experiment, we assayed plant biomass based on estimates of leaf surface area by taking standardized digital images which were analyzed with ImageJ software and a two-tailed Student’s *t*-test.

### Soil microbial activity

We measured respiration of the soil substrate on a weekly basis to determine the effect of Rpf treatment on belowground microbial activity. For each pot, we transferred 1 g of soil substrate into glass vials with a silicon membrane septum cap. After incubating at 25 °C in the dark for 24 h, we measured the CO_2_ concentration (ppm) from 1 mL of headspace gas using an infrared gas analyzer (IRGA) (LI-COR Environmental). We then estimated CO_2_ concentration in our samples based on values generated from a standard curve of known CO_2_ concentrations. We performed repeated-measures ANOVA (RM-ANOVA) using an AR(1) covariance structure to test for the main effects and interaction of Rpf and time on rates of respiration.

### Microbial abundance

We estimated bacterial and fungal abundance using quantitative PCR (qPCR) on DNA extracted from soil substrate. For each pot, we collected 1 g of soil substrate pooled from three subsamples and immediately stored them at −80 °C. We extracted the DNA from the sample using a PowerSoil® Total RNA Isolation Kit with DNA Elution Accessory Kit (MoBio) following manufacturer’s protocol, and the DNA was subsequently quantified using a Take5 *Synergy* microplate reader (BioTek). We then conducted qPCR assays described in greater detail elsewhere (Fierer, Jackson, Vilgalys, & Jackson, 2005; Lau & Lennon, 2011). Briefly, each 20 μL reaction contained 1 μL of DNA template (2.5 ng μL^-1^), 1 μL of each primer (10 μmol L^-1^), 7 μL of molecular-grade water, and 10 μL iQTM SYBR Green SuperMix (Bio-Rad Laboratories Hercules, CA, USA). For bacteria we used the Eub338 forward primer (ACTCCTACGGGAGGCAGCAG) (Lane, 1991) and the Eub518 reverse primer (ATTACCGCGGCTGCTGG) (Muyzer, Dewaal, & Uitterlinden, 1993) to amplify the 16S rRNA gene, and for fungi we used the ITS1f forward primer (TCCGTAGGTGAACCTGCGG) (Gardes & Bruns, 1993) and the 5.8S reverse primer (CGCTGCGTTCTTCATCG) (Vilgalys & Hester, 1990) to amplify the ITS gene region. qPCR assays were performed using Eppendorf Mastercycler Realplex system using the previously reported thermal cycle conditions: 15 min at 95 °C, followed by 40 cycles of 95 °C for 1 min, 30 s at 53 °C for annealing, followed by 72 °C for 1 min (Fierer et al., 2005). The coefficients of determination (*r*^2^) of our assay ranged from 0.95 and 1, while amplification efficiencies fell between 0.93 and 0.99. Based on melting curve analyses, we found no evidence for primer dimers. We estimated fungal and bacterial abundance based on the estimated gene copy number from their respective standard curves generated from bacterial and fungal isolates as described elsewhere (Lau & Lennon, 2012). We performed a twotailed Student’s *t*-test to determine the effect of the Rpf treatment on soil bacterial (16S rRNA) and fungal (ITS) gene copy abundances, as well as the fungal to bacterial ratio (F: B).

### Microbial diversity

To account for the variation in metabolic activity, we characterized bacterial communities from the soil substrate using pools of RNA and DNA. DNA is a relatively stable molecule contained in intact cells irrespective of their metabolic status. Accordingly, we interpret 16S rRNA sequences recovered from the DNA pool as the “total” community. In contrast, RNA is a more ephemeral molecule that is required for protein synthesis by actively growing cells. Therefore, we interpret 16S rRNA sequences recovered from the RNA pool after complementary DNA (cDNA) synthesis as the “active” community (Jones & Lennon, 2010). We assume that dormant individuals can create discrepancies between the active and total composition of a given sample, but we do not attempt to use RNA and DNA to directly characterize dormant taxa. Following previously described protocols (Lennon, Placella, & Muscarella, 2017), we characterized the effects of Rpf on soil bacterial communities using high-throughput sequencing. First, we extracted nucleic acids using the PowerSoil Total RNA Extraction Kit with DNA Elution Accessory Kit (MoBio, Carlsbad, CA) followed by cleaning via ethanol precipitation. We removed residual DNA from RNA samples using DNase 1 (Invitrogen) following manufacturer’s protocol and synthesized cDNA by reverse transcribing RNA using iScript Reverse Transcription Supermix for RT-qPCR (Bio-Rad). After ensuring that there was no product in our no-template negative controls, we amplified the V4 region of the 16S rRNA gene from bacterial DNA and cDNA using the 515F primer (GTGYCAGCMGCCGCGGTAA) and the 806R primer (GGACTACNVGGGTWTCTAAT) that included a unique barcode for each sample (Caporaso et al., 2012). Thermal cycle conditions for the PCR reaction consisted of 3 min at 94 °C, followed by 30 cycles of 94 °C for 45 s, 30 s at 50 °C, 90 s at 72 °C, followed by 72 °C for 10 min. We cleaned the sequence libraries using the Agencourt AMPure XP purification kit (Beckman Coulter, Brea, CA, USA), quantified the resulting products using the QuanIt PicoGreen kit (Invitrogen), and pooled libraries at equal molar ratios (final concentration: 10 ng per library). We then sequenced the pooled libraries with the Illumina MiSeq platform using 250 x 250 bp paired end reads (Illumina Reagent Kit v2, 500 reaction kit) at the IU-CGB. Paired-end raw 16S rRNA sequence reads were assembled into contigs using the Needleman algorithm (Needleman & Wunsch, 1970). We then trimmed the resulting sequences using a moving average quality score (window = 50 bp, score = 35), in addition to removing long sequences, ambiguous base calls, and sequences that matched Archaea, chloroplasts, and other non-bacteria. We also removed chimeric sequences that were detected using the UCHIME algorithm (Edgar, Haas, Clemente, Quince, & Knight, 2011). After this filtering, there was a total of 3,217,419 quality reads. We aligned the sequences to the Silva Database (v 128) using the Needleman algorithm (Needleman & Wunsch, 1970). We created operational taxonomic units (OTUs) by first splitting the sequences based on taxonomic class (using the RDP taxonomy) and then binning sequences into OTUs based on 97 % sequence similarity using the OptiClust algorithm (Westcott & Schloss, 2017). We removed all OTUs with less than two occurrences in the data set. Altogether, this led to a high degree of coverage across samples (minimum Good’s coverage = 0.97). All sequence processing was completed using the software package *mothur* (version 1.39.5) (Schloss et al., 2009).

First, we used the 16S rRNA sequences to test how the Rpf treatment affected measures of alpha diversity with samples. We estimated bacterial richness (i.e., the number of OTUs in a sample) using a resampling with replacement approach that subsampled 1,000 sequence observations per sample, resampled 999 additional times, and then calculated the average number of OTUs to estimate per sample richness (± SEM) (Gotelli & Colwell, 2011; Muscarella, Jones, & Lennon, 2016). Similarly, we estimated the effect of Rpf treatment on taxa evenness (i.e., the equitability in relative abundance of OTUs in a sample) using the same resampling approach with the Evar index (Smith & Wilson, 1996). We tested for the effect of Rpf on these measures of diversity using ANOVA.

Second, we used the sequence data to test how the Rpf treatment affected beta diversity among samples. Given that relative abundances spanned order of magnitudes, we log10-transformed data to prevent undue weight of dominant species. Following this, we conducted Principal Coordinates Analyses (PCoA) using the Bray-Curtis dissimilarity metric to visualize the effects of Rpf on bacterial communities. To test the hypothesis that Rpf affected the composition of the total (DNA) and active (RNA) bacterial pools, we used permutational multivariate analysis of variance (PERMANOVA) with Bray-Curtis dissimilarity metric implemented with the `adonis` function in the *vegan* package (Oksanen et al., 2011) in the R statistics environment (v 3.2.3). We coded pot number as a factor to account for paired-sample design given that DNA and RNA were co-extracted from the same soil-substrate sample within an experimental unit. Last, we conducted indicator species analyses to identify influential taxa driving compositional changes in response to the Rpf treatment. Specifically, we calculated Pearson’s phi coefficients of association with taxa abundance data using the `multipatt` function in the *indicspecies* R package (De Cáceres & Legendre, 2009). We filtered the output from the indicator species analyses so as to only consider associations between taxa and treatments where *P*-values were ≤ 0.05 and correlations were ≥ |0.7|.

## RESULTS

### Plant responses

Plants in +Rpf treatment had less total (*t*_7_ = 2.84, *P* = 0.013), aboveground (*t*_7_ = 2.70, *p* = 0.017), and belowground biomass (*t*_7_ = 2.30, *P* = 0.049) than plants in the -Rpf treatment (Fig. 2). While recombinant protein led to a ~33 % reduction in root biomass, Rpf had no effect on other plant traits including SLA (*t*_7_ = 0.80, *P* = 0.442), shoot: root ratio (*t*_7_ = −1.20, *P* = 0.250), shoot height (*t*_7_ = 1.11, *P* = 0.286), or the number of seeds produced per plant (*t*_7_ = 0.83, *P* = 0.421) (Fig. S2). However, we did detect a marginal decrease in the number of flowers produced per plant in the +Rpf treatment (*t*_7_ = 1.79, *P* = 0.097). In the *Arabidopsis* experiment where individuals were grown in the absence of soil-substrate and an accompanying microbiome, there was no significant effect of Rpf on estimates of plant biomass (*t*_3_ = −1.11, *P* = 0.320, Fig. S3).

**Fig. 2.**
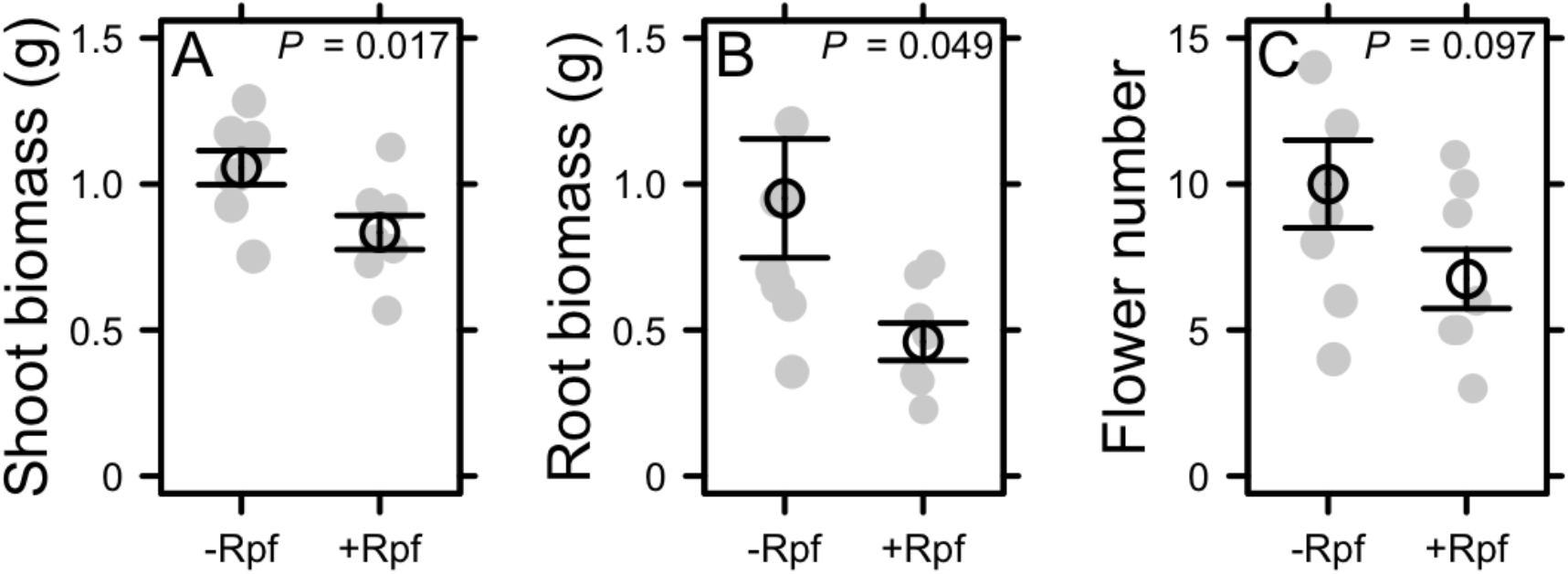
Influence of resuscitation promoting factor (Rpf) on (A) *Brassica rapa* shoot biomass, (B) root biomass, and (C) flower number produced per plant. We compared plant traits from individuals that were exposed to weekly additions of recombinant Rpf (+Rpf) to those exposed to a protein buffer control (-Rpf). Black symbols represent the mean ± 95 % confidence intervals. Grey symbols represent the individual observations.

### Microbial activity

Rpf altered the activity of the belowground microbial community based on respiration of the soil substrate. We detected a significant main effect of Rpf (*F*_1, 84_ = 9.68, *P* = 0.002) and time (*F*_5, 84_ = 21.60, *P*< 0.001), but no interaction (*F*_5, 84_ = 1.22, *P* = 0.308), on respiration (Fig. 3). While respiration increased throughout the experiment in both treatments, based on marginal means, soil respiration was 24 % lower in the +Rpf treatment than in the -Rpf treatment.

**Fig. 3.**
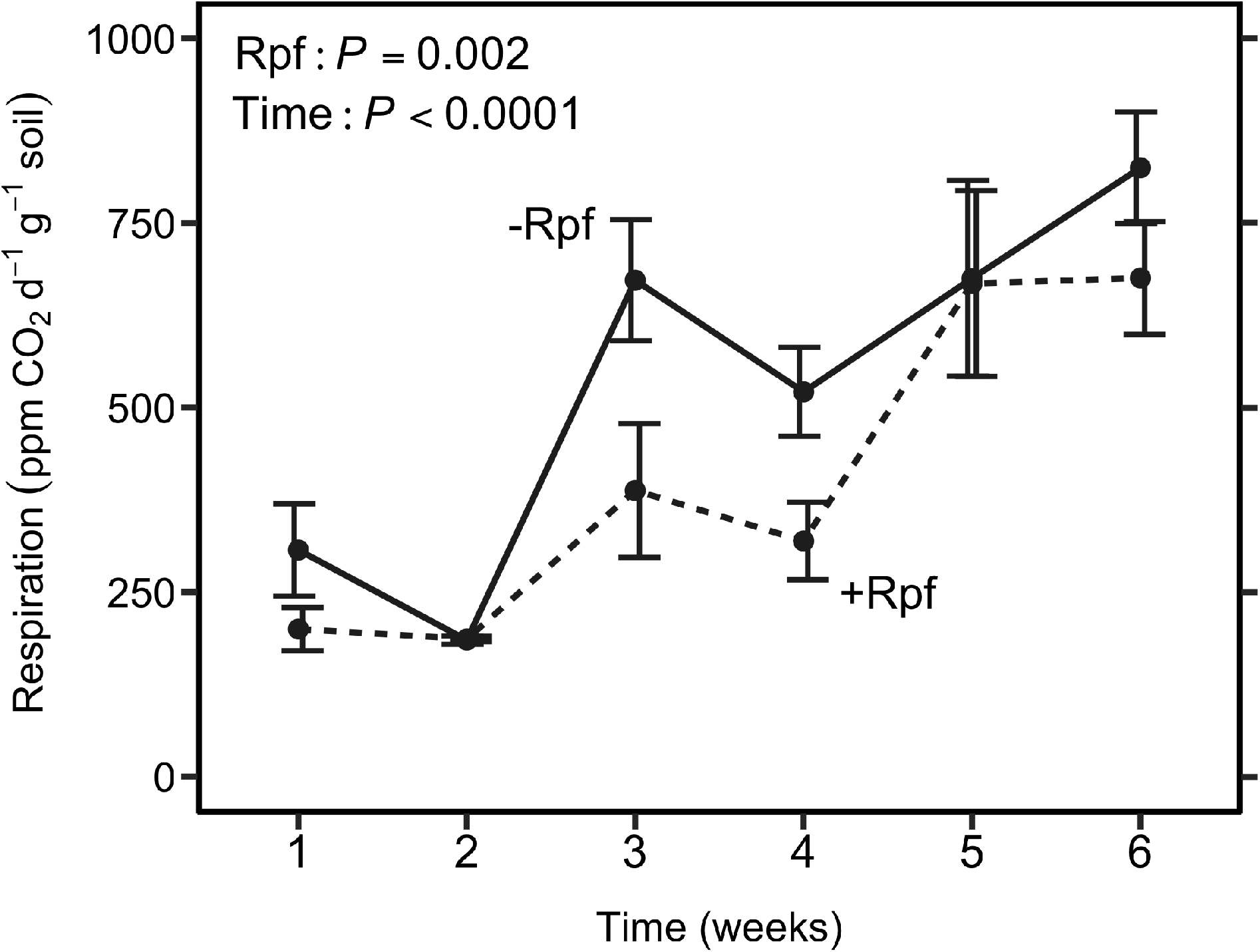
Effect of resuscitation promoting factor (Rpf) on soil microbial activity. Soil respiration was measured after applying Rpf (+Rpf) or protein buffer control (-Rpf) to soils on a weekly basis. Symbols represent the mean ± 1 SE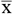.

### Microbial abundance

The Rpf treatment significantly altered the abundance of soil bacteria (*t*_8_ = 2.71, *P* = 0.016) and fungi (*t*_8_ = −3.27, *P* = 0.007) relative to the control (Fig. 4). Bacterial abundance estimated as 16S rRNA gene copy number decreased by ~30 % in the +Rpf treatment relative to the -Rpf treatment. In contrast, fungal abundance estimated as the ITS gene copy number increased 2.8-fold in the +Rpf treatment. Consequently, soil F: B ratio increased 4.8-fold under +Rpf treatment relative to the control treatment (*t*_8_ = −2.84, *P* = 0.018).

**Fig. 4.**
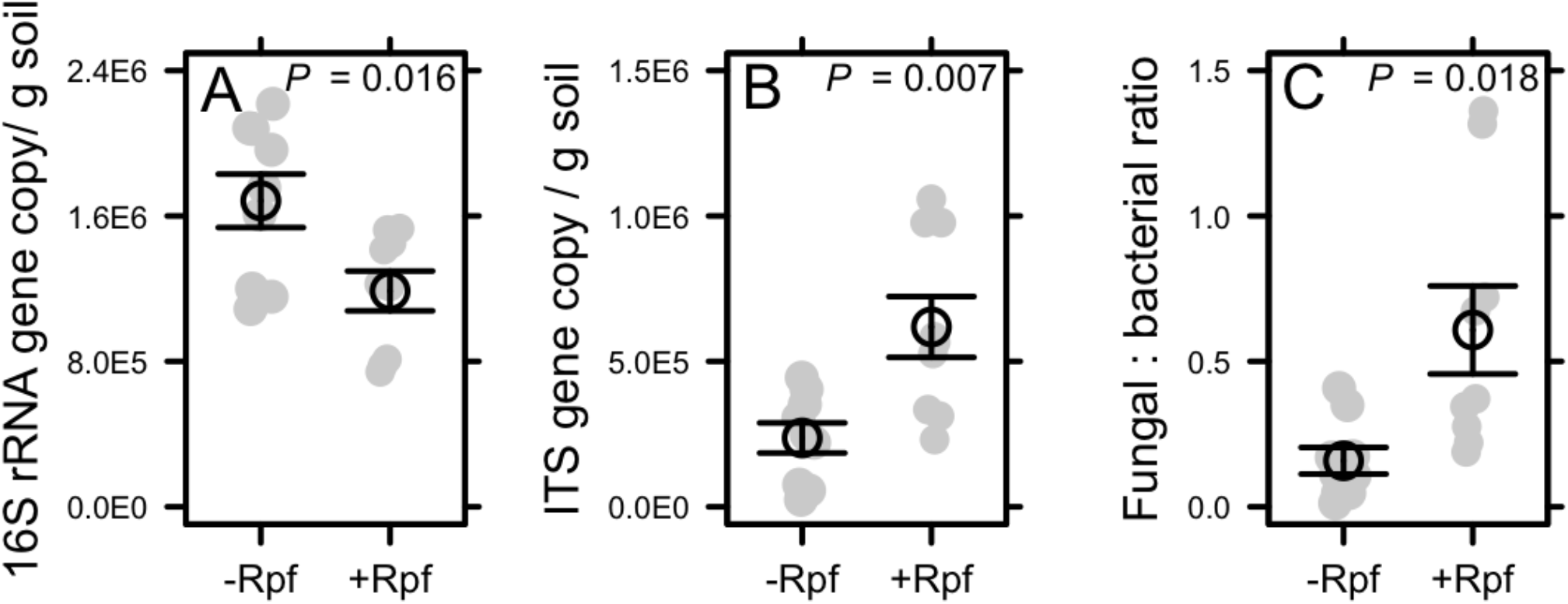
Effect of resuscitation promoting factor (Rpf) on (A) soil bacterial 16S rRNA copy number, (B) fungal ITS copy number, and (C) the fungi: bacteria gene copy ratio. Gene copy number was measured from soil after six weekly applications of recombinant Rpf (+Rpf) or protein buffer control (-Rpf) treatment. Black symbols represent the mean ± 95 confidence intervals. Grey symbols represent the individual observations.

### Bacterial diversity

Bacteria in the soil-substrate were diverse and reflected compositions that were not unlike many natural communities. Based on 16S rRNA sequencing, we recovered a total 32,055 taxa across the experimental units. The bacterial community was dominated by OTUs belonging to the following phyla: Proteobacteria (49 %), Acidobacteria (14 %), Verrucomicrobia (7 %), Planctomycetes (7 %), Bacteroidetes (5 %), Actinobacteria (4 %), Chlorofexi (2 %), and Firmicutes (2 %).

To evaluate broad-scale treatment effects on microbial diversity, we first examined changes among the major groups of bacteria. Rpf additions had no effect on bacterial richness regardless of whether sequences came from the total (*F*_1, 8_ = 0.44, *P* = 0.526) or active community (*F*_1, 8_ = 0.047, *P* = 0.835) (Table S2). Similarly, Rpf had no effect on the evenness of the total (*F*_1, 8_ = 0.527, *P* = 0.489) or active (*F*_1, 8_ = 0.108, *P* = 0.751) community (Table S2). In contrast, bacterial composition was significantly affected by metabolic status (i.e., active vs total) (*F*_1, 19_ = 7.24, *r*^2^ = 0.23, *P* = 0.001) and the Rpf treatment (*F*_1, 19_ = 1.5, *r*^2^ = 0.06, *P* = 0.032), which can be visualized in Fig. 5. Indicator species analysis revealed that there were 137 taxa belonging to the Proteobacteria and Bacteroidetes that were associated with the + Rpf treatment. Meanwhile, 195 taxa in the Planctomycetes, Proteobacteria, Chlorflexi, Acidobacteria, and Actinobacteria were significantly associated with the -Rpf treatment.

**Fig. 5.**
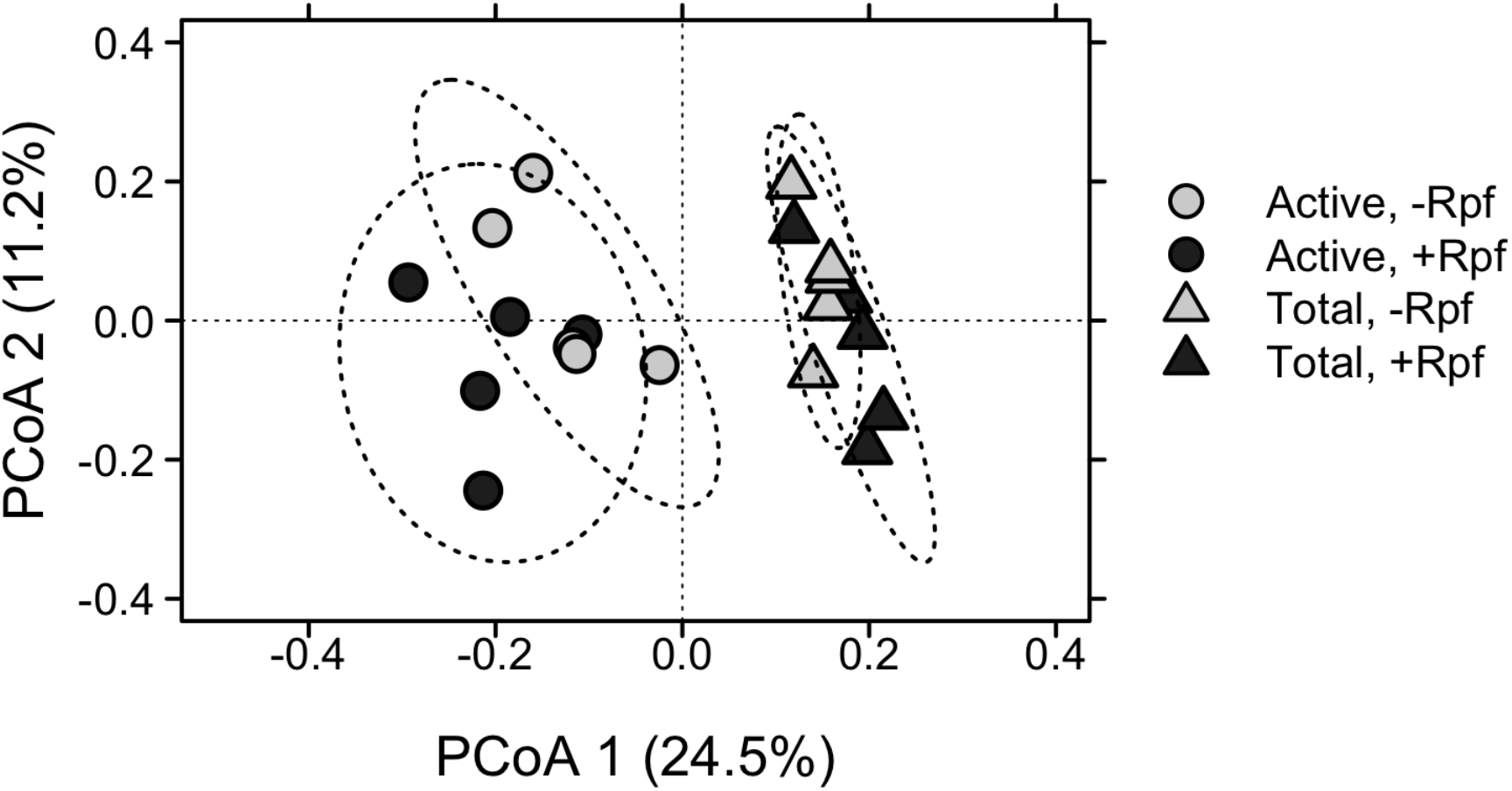
Principal Coordinate Analysis (PCoA) plot depicting composition of active (i.e., RNA) and total (i.e., DNA) Actinobacteria from soil that were exposed to +Rpf and -Rpf treatments at the end of a six-week experiment. The ellipses were generated by ‘ordiellipse’ function using the standard deviation of PCoA point scores to visualize the spread of each treatment.

Since the production of Rpf is thought to be restricted to the Actinobacteria, we also evaluated how treatments affected the diversity of taxa within this phylum. Rpf had no effect on the richness (total community: *F*_1, 8_ = 0.80, *P* = 0.397; active community: *F*_1, 8_ = 0.14, *P* = 0.722), nor did it influence evenness (total community: *F*_1, 8_ = 0.69, *P* = 0.429; active community: *F*_1, 8_ = 0.41, *P* = 0.538) (Table S3). There was a small and marginally significant reduction in the relative abundance of actinobacterial sequences in the +Rpf treatment compared to the -Rpf treatment (*t*_9_ = 1.46, *P* = 0.083) (Fig. S4). Based on PERMANOVA, actinobacterial composition was significantly affected by both metabolic status (i.e., active vs total) (*F*_1, 19_ = 16.08, *r*^2^ = 0.33, *P* = 0.001) and the Rpf treatment (*F*_1, 19_ = 3.25, *r*^2^ = 0.07, *p* = 0.010) (Fig. S5). Indicator species analysis revealed that OTUs belonging to the *Acidothermus, Catenulispora*, and *Mycobacterium* were associated with the -Rpf treatment, while only a single taxon belonging to the Solirubrobacterales had a significant association with the +Rpf treatment (Fig. S6)

## DISCUSSION

Dormancy is an important life-history trait that influences the diversity, composition, and function of soil microbial communities. After generating recombinant protein with a gene from an environmental isolate, we applied resuscitation promoting factor (Rpf) to a soil community to test the hypothesis that microbial seed-bank dynamics alter plant growth and fitness traits by driving changes in the belowground microbial community (Fig. 1). The Rpf treatment decreased plant biomass (Fig. 2) most likely by altering the activity (Fig. 3), abundance (Fig. 4), and composition (Fig. 5) of soil microbial communities. These findings are consistent with the view that soil microbial seed banks can influence plant performance, perhaps by disrupting interactions with beneficial microorganisms or through the recruitment of pathogens. In the following sections, we discuss these findings while exploring other potential ways in which Rpf may affect plant-microbe interactions in soil environments.

### Rpf indirectly affects plant performance

Resuscitation of microbial seed banks led to a 33 % reduction in plant biomass (Fig. 2). For the following reasons, we infer that Rpf effects on plant performance are likely to be indirect. First, it is known that Rpf activity involves the hydrolysis of glyosidic linkages in peptidoglycan (Cohen-Gonsaud et al., 2005; Mukamolova et al., 2006).While this polymer is found in the cell walls of virtually all bacteria, it is absent from plant tissue. Second, we conducted an experiment with *Arabidopsis thaliana*, a fairly close relative of *B. rapa*, to evaluate whether Rpf can directly affect plant performance. We detected no significant treatment effect after maintaining a relatively small number of seedlings on microbe-free agar for five weeks. However, there was a trend of reduced plant growth (18 %) for individuals in the +Rpf treatment. While additional experiments may be warranted, for the purposes of this study, we cautiously conclude that Rpf influenced plants primarily via its effect on the activity, abundance, or composition of the soil microbial community.

### Rpf altered fungal-bacterial interactions

Rpf may have altered plant performance by modifying fungal-bacterial interactions. We observed a nearly three-fold increase in fungal abundance in response to Rpf additions (Fig. 4). As a result, soil F: B ratio increased under Rpf treatment (Fig. 4), which may also explain the reduction in soil microbial activity (Fig. 3) given that there are often differences in carbon-use efficiency between bacteria and fungi (Sakamoto & Oba, 1994; Whitaker et al., 2014). Shifts in F: B may also reflect the way that Rpf influences microorganisms with different cell-wall properties. Like plants, fungi do not contain peptidoglycan, the substrate which Rpf acts upon. Instead, the fungal cell wall is primarily composed of chitin. Therefore, an increase in fungal abundance could arise if Rpf decreased the competitive ability of soil bacteria in our system. In principle, the muralytic activity of Rpf may have even lysed some bacterial cells. If this occurred, the resulting necromass could be scavenged by fungi to meet their metabolic demands (Bradley, Amend, & LaRowe, 2018). Our results, however, do not support this hypothesis. We found that bacterial abundance from three contrasting soils increased with Rpf concentration up to 4 μM g^-1^ soil. At higher concentrations (8 μM g^-1^), we no longer observed a growth-stimulating effect, but bacterial abundance never dropped below levels observed in control soils without Rpf (Fig. S6). Rpf was much lower (40 nM g^-1^ soil) in our *Brassica* experiment, yet above concentrations shown to resuscitate dormant bacteria (Cohen-Gonsaud et al., 2005; Mukamolova et al., 1998). In addition to demonstrating the generality of our recombinant protein on different soils, these findings suggest that it is unlikely that reductions in bacterial abundance (Fig. 4) were due to lysis or direct inhibition of the soil microbial community.

### Rpf altered microbial diversity

While seed banks are known to promote biodiversity by buffering ecological and evolutionary dynamics, microbial dormancy may have complex effects that influence plant-soil dynamics. The influence of Rpf on communities of interacting species is likely dependent on enzyme specificity. Very few studies have investigated Rpf effects on a broad range of microorganisms, but some evidence suggests cross-species resuscitation using recombinant Rpf generated from actinobacterial strains belonging to *Mycobacterium* and *Brevibacterium* (Mukamolova et al., 1998; Puspita et al., 2015).

In our study, we found that Rpf had broad-scale effects on bacterial communities. Results from an indicator species analysis suggest that some taxa may have been inhibited by the treatment, while others belonging to the Proteobacteria and Bacteroidetes were favored by Rpf additions. These findings imply that Rpf from a single species may resuscitate a wide range of taxa, consistent with reports of a conserved catalytic site within the peptidoglycan substrate (Mukamolova et al., 2002). Alternatively, Rpf may only stimulate growth in a smaller set of taxa, but their metabolism leads to a cascade of community change. Our indicator species point to one such group of taxa within the Actinobacteria, which are the only known producers of Rpf. Specifically, OTUs belonging to the *Solirubrobacterales* were associated with the +Rpf treatment. Members of this group are aerobic, non-spore-forming, and non-motile bacteria that are often recovered in forest and agricultural soils (Kim et al., 2007; Reddy & Garcia-Pichel, 2009; Seki, Matsumoto, Ōmura, & Takahashi, 2015). Little is known about the role of *Solirubrobacterales* in the context of plant-microbe interactions, but some representatives assimilate simple organic compounds (e.g., glucose, maltose, sucrose, xylose, and arginine) that are commonly found in the rhizosphere (Reddy & Garcia-Pichel, 2009). Interestingly, the *Solirubrobacterales* are only distantly related to *Micrococcus* KBS0714, the soil isolate containing the Rpf gene that we used for producing the purified recombinant protein. Again, this suggests that Rpf functionality may not be restricted to close kin. Additional work is needed to unveil the network of interactions among Rpf producers and responders, in addition to whether or not Rpf genes map onto the phylogeny of Actinobacteria.

### Implications of seed-bank dynamics for plant-microbe interactions

Our study demonstrates that resuscitation promoting factor (Rpf) modified plant performance most likely through its effects on soil microbial communities. Reductions in plant growth were accompanied by shifts in soil microbial community properties, which may have included the recruitment of pathogens. Although we did not observe root lesions or other clear signs of infection, we cannot rule out the possibility the seed banks harbor pathogenic or non-mutualistic taxa that could decrease plant performance. Such a view is consistent with reports of microbial pathogens persisting in soil for long periods of time, likely in a state of reduced metabolic activity (Sharma & Reynnells, 2016). More explicit tests would include the targeting of known pathogens combined perhaps with measurements of plant immune responses to altered microbial seed banks. Alternatively, our findings could be explained by a dilution of mutualists in the soil or a general disruption of beneficial plant-microbe interactions, which would likely be reflected in the profiling of plant exudates and microbial metabolites.

While our study provides empirical support for the notion that microbial seed banks can affect plant performance, it also raises a number of questions for future exploration. We observed that a one protein from a single strain of bacteria can have broad effects on interacting plants and microorganisms. However, some bacteria may be responsive to certain families of Rpf, while others may not. This likely is due to the mechanisms by which Rpf operates in a community context, which is a topic that has received little attention to date. More generally, metabolic transitions into and out of dormancy are influenced by mechanisms besides Rpf (Lennon & Jones, 2011) many of which are tied to the interpretation of environmental triggers (e.g., soil rewetting) that are known to resuscitate microorganisms (Aanderud, Jones, Fierer, & Lennon, 2015). Finally, there is a need to scale up manipulative experiments to explore microbial seed banks in more complex communities. Generalities could be assessed by examining the effects of Rpf outside of the Brassicaceae given that these plants tend to lack mycorrhizal symbionts, which are important mutualists found among most flowering plants. It is not difficult to imagine that more complex outcomes could emerge when altered microbial seed banks are considered in diverse plant communities (Fig. 1). Nevertheless, our study highlights a potentially important but overlooked component of plant-microbe interactions. It is estimated that soil microbial communities can be dominated by dormant taxa, yet changes in activity can be fast, suggesting that seed banks may be an important factor contributing to plant-soil feedbacks, which is thought to be an important mechanism maintaining landscape patterns of biodiversity (Bever, 2003).

## ACKNOWLEDGEMENTS

We acknowledge Peyton Thomas for assistance in the laboratory and members of the Lennon lab for critical feedback on an earlier version of the manuscript. This work was supported by the National Science Foundation (1442246 JTL, 1934554 JTL), the US Army Research Office Grant (W911NF-14-1-0411 JTL), and the National Aeronautics and Space Administration (80NSSC20K0618 JTL).

## AUTHOR CONTRIBUTIONS

V.K. and J.T.L. designed the study; V.K. and B.K.L performed the experiments; V.K. and J.T.L analyzed the data; V.K. and J.T.L. wrote the paper.

## DATA AVAILABILITY STATEMENT

All DNA sequences can be downloaded from the NCBI BioProject PRJNA504042. Other data and code are available from the Dryad Digital Repository (https://doi.org/10.XXX/XXXXX) and GitHub (https://github.com/LennonLab/BrassicaRpf)

