## supplemental for "Resuscitation of the microbial seed bank alters plant-soil interactions"

**Table 1.** Characteristics of surface soil (0- 5 cm) from Griffy Woods at the Indiana University Research and Teaching Preserve. These soils were mixed with equal volume of metro-mix and vermiculite to create a suitable substrate for *Brassica rapa* in growth chambers. Analyses were performed by Ward Laboratories, Inc. (https://www.wardlab.com/).

| **Variable** | **Value** |
|---|---|
| pH | 5.5 |
| Soluble salts | 0.09 mS cm-1 |
| Organic matter | 5.2 % |
| Nitrate-N | 0.4 ppm |
| Phosphorus | 7 ppm |
| Potassium | 130 ppm |
| Sulfate | 12.1 ppm |
| Calcium | 738 ppm |
| Magnesium | 124 ppm |
| Sodium | 124 ppm |
| Nitrogen, Total | 2029 ppm |
| Carbon, Total | 2.7 % |
| Sum of cation | 9.3 meq100g soil-1 |
| % Base saturation | |
| H+ | 45 |
| K+ | 4 |
| Ca+ | 39 |
| Mg+ | 11 |
| Na+ | 1 |
| Soil texture | |
| Sand | 30% |
| Silt | 56% |
| Clay | 14% |

**Table S2**. Taxon richness and evenness (i.e., alpha diversity) diversity of soil bacteria in the two Rpf treatments (-Rpf and +Rpf) for total (DNA) and active (RNA) pools ($n$ = 5). We calculated richness as the number of operational taxonomic units (97 % sequence similarity of the 16S rRNA gene) and evenness using Smith and Wilson's Evenness Index (Evar). Values are means and standard error of the means (in parentheses). There was no significant effect of Rpf treatment on soil bacterial richness in the total ($F_{1, 8}$ = 0.44, $P$ = 0.526) or active pool ($F_{1, 8}$ = 0.047, $P$ = 0.835). Likewise, Rpf treatment did not alter the bacterial community evenness in the total ($F_{1, 8}$ = 0.527, $P$ = 0.489) or active pool ($F_{1, 8}$ = 0.108, $P$ = 0.751).

| Treatment | Richness | | Evenness | |
|---|---|---|---|---|
| | Active | Total | Active | Total |
| -Rpf | 481 | 498 | 0.73 | 0.75 |
| | (24.3) | (16.6) | (0.175) | (0.013) |
| +Rpf | 488 | 475 | 0.74 | 0.732 |
| | (20.8) | (30.7) | (0.015) | (0.022) |

**Table S3**. Taxon richness and evenness (i.e., alpha diversity) diversity of soil Actinobacteria in the two Rpf treatments (-Rpf and +Rpf) for total (DNA) and active (RNA) pools ($n$ = 5). We calculated richness as the number of operational taxonomic units (97 % sequence similarity of the 16S rRNA gene) and evenness using Smith and Wilson's Evenness Index (Evar). Values are means and standard error of the means (in parentheses). There was no significant effect of Rpf treatment on soil actinobacterial richness in the total ($F_{1,8} = 0.68$, $P = 0.435$) or active pool ($F_{1,8} = 0.01$, $P = 0.923$). Likewise, Rpf treatment did not alter the actinobacterial community evenness in the total ($F_{1,8} = 0.245$, $P = 0.634$) or active pool ($F_{1,8} = 0.005$, $P = 0.944$).

| Treatment | Richness | | Evenness | |
|---|---|---|---|---|
| | Active | Total | Active | Total |
| -Rpf | 155 | 171 | 0.442 | 0.457 |
| | (17.0) | (4.6) | (0.021) | (0.004) |
| +Rpf | 157 | 164 | 0.444 | 0.453 |
| | (8.7) | (6.1) | (0.013) | (0.007) |

**Fig. S1.** Western blot confirming the presence and expected size of the recombinant Rpf protein that was purified and eluted using the Ni-NTA Purification System (Invitrogen). We performed the Western blot using the XCell *SureLock*™ Mini-Cell (Invitrogen) with a NuPAGE Bis-Tris Mini Gel (Novex) to run the electrophoresis gel according to the manufacturer's instructions. Next, we performed the transfer run using the XCell II™ Blot Module (Invitrogen) with a 0.2 µm pore size PVDF membrane (Invitrogen) according to the manufacturer's instructions. Briefly, 5 µl of recombinant Rpf protein (2.6 mg/mL) was diluted ten-fold, heat-denatured, and run on gel electrophoresis immersed in 1X running buffer (Invitrogen) following recommended run conditions. Afterwards, we performed a transfer run using 1X transfer buffer (Invitrogen) using PVDF membrane filter paper following recommended run conditions. We then performed membrane blocking at 22 °C with gentle rocking following these conditions: an initial blocking for 10 m with 5 mL of StartingBlock (ThermoFisher), incubation with primary IgG anti-Rpf serum (1:50000) for 2 h, 4X membrane washing with 25 mL TBST for 10 m each, incubation with secondary IgG anti-rabbit (alkaline phosphatase) (1:20000), and 4X membrane washing with 25 mL of TBST for 10 m each. Finally, we added 1 mL of detection reagent 1-Step NBT/BCIP to the immunoblotted membrane to generate visible protein banks resulting from the conjugated antibodies reacting with the detection dye. Once protein bands appeared, the membrane was immediately immersed in DI water to stop reaction. Our final Western blot membrane shows purple bands depicting 5 and 10 µl of pET15b recombinant Rpf at 900 µg/mL concentration bound with anti-Histidine (1:50000) primary IgG in lane 2 and 4, respectively. Lane 1 and 3 are empty. Lane 5 is a low-range Western marker (red/40, blue/15, green/10, blue/2.6, blue/1.7 kDa). The Western blot clearly shows the presence of Rpf protein eluted from the Ni-NTA column that is ~40 kDa in size.

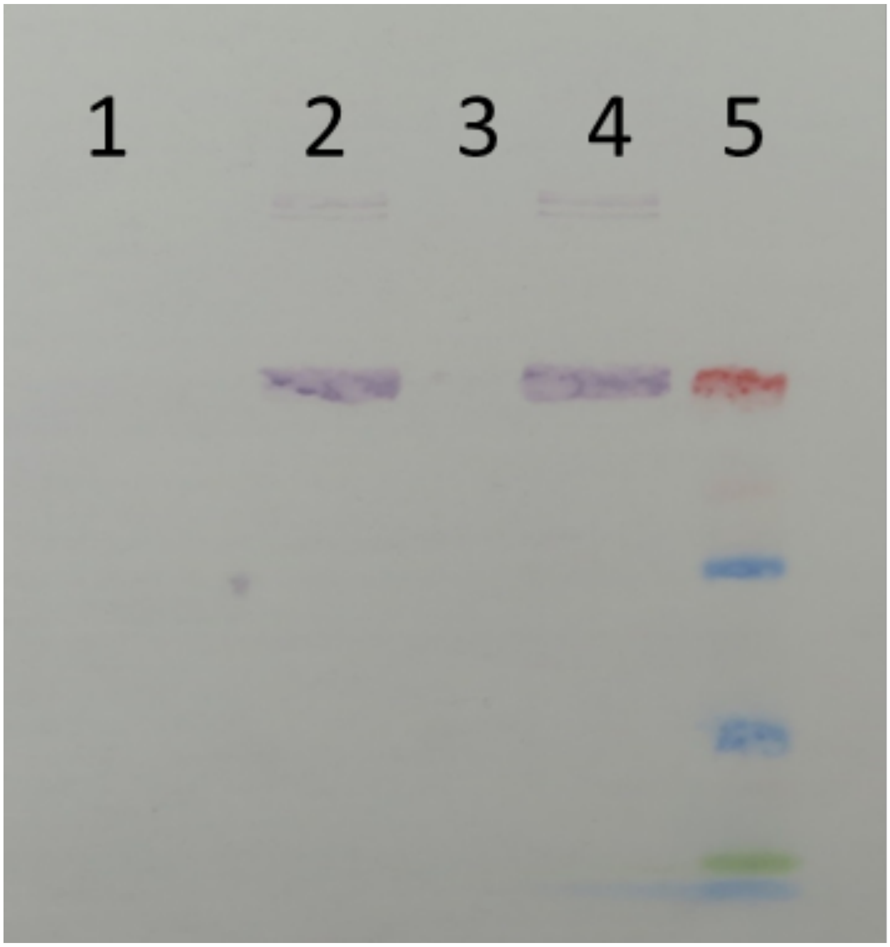
1
2
3
4
5

**Fig. S2.** Influence of resuscitation promoting factor (Rpf) on *Brassica rapa* (A) shoot : root ratio, (B) shoot height (cm), (C) seed production, and (D) specific leaf area. We compared plant traits from individuals that were exposed to weekly additions of recombinant Rpf (+Rpf) to those exposed to a protein buffer control (-Rpf). Black symbols represent the mean ± 95 % confidence intervals. Grey symbols represent the individual observations.

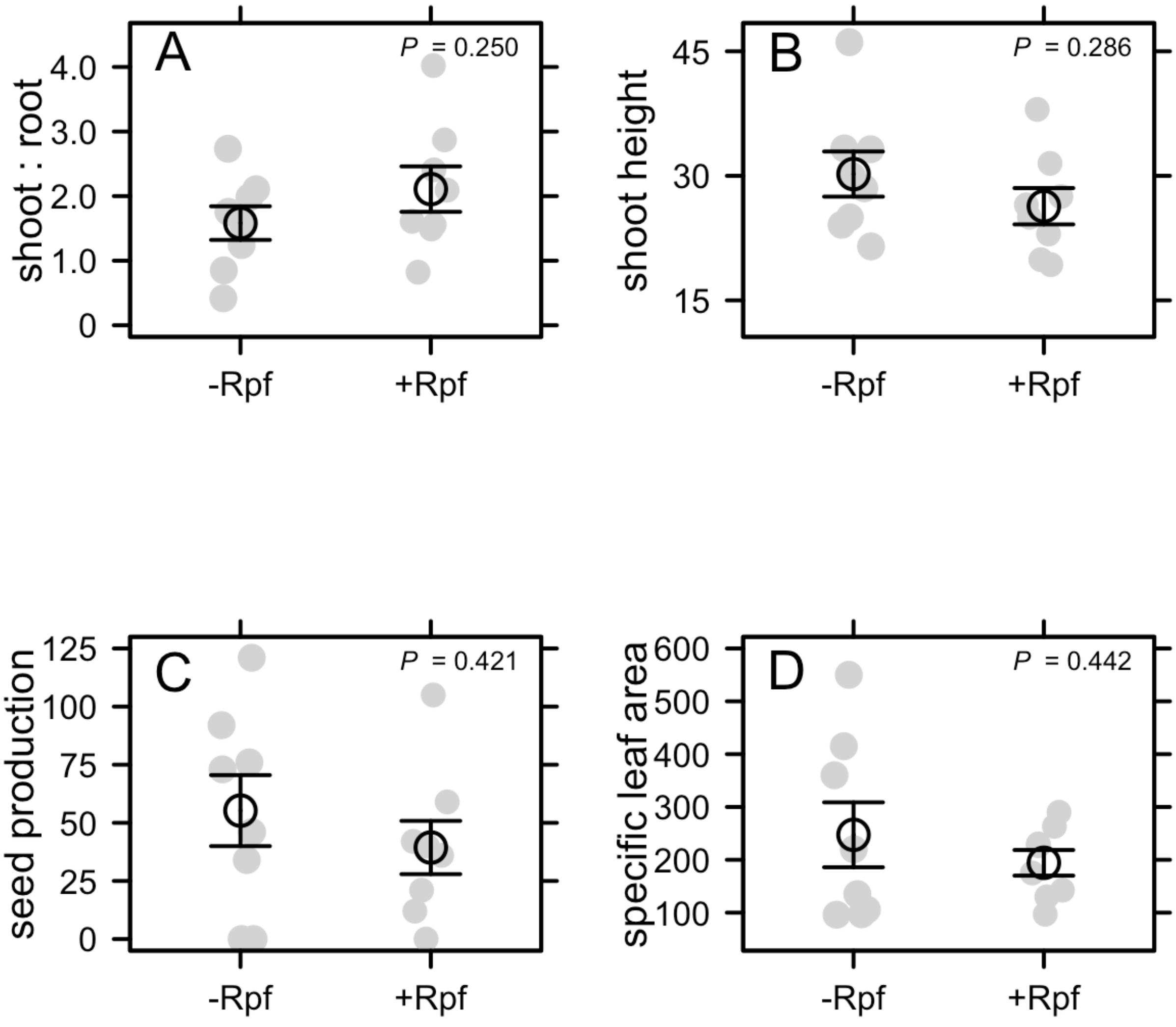

**Fig. S3**. Effects of Rpf on *Arabidopsis thaliana* seedlings grown on sterile Murashige-Skoog (MS) agar plates exposed to either Rpf protein (final concentration: 1.6 µmol/L) (+Rpf) or protein buffer control (-Rpf) after five weeks. One replicate in the -Rpf treatment was excluded due to microbial contamination. Black symbols represent mean ± 95 % confidence intervals. Grey symbols represent individual observations.

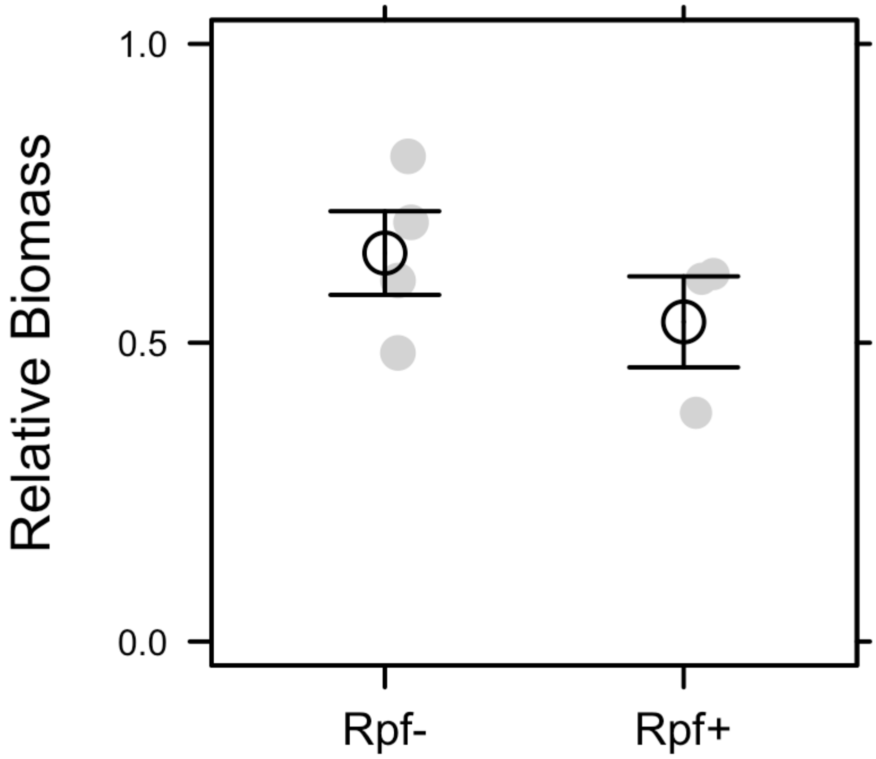

**Fig. S4**. The effect of resuscitation promoting factor (Rpf) on the relative abundance of Actinobacteria to all other 16S rRNA bacterial sequences at the end of the experiment. Black symbols represent mean ± 95 % confidence intervals. Grey symbols represent individual observations.

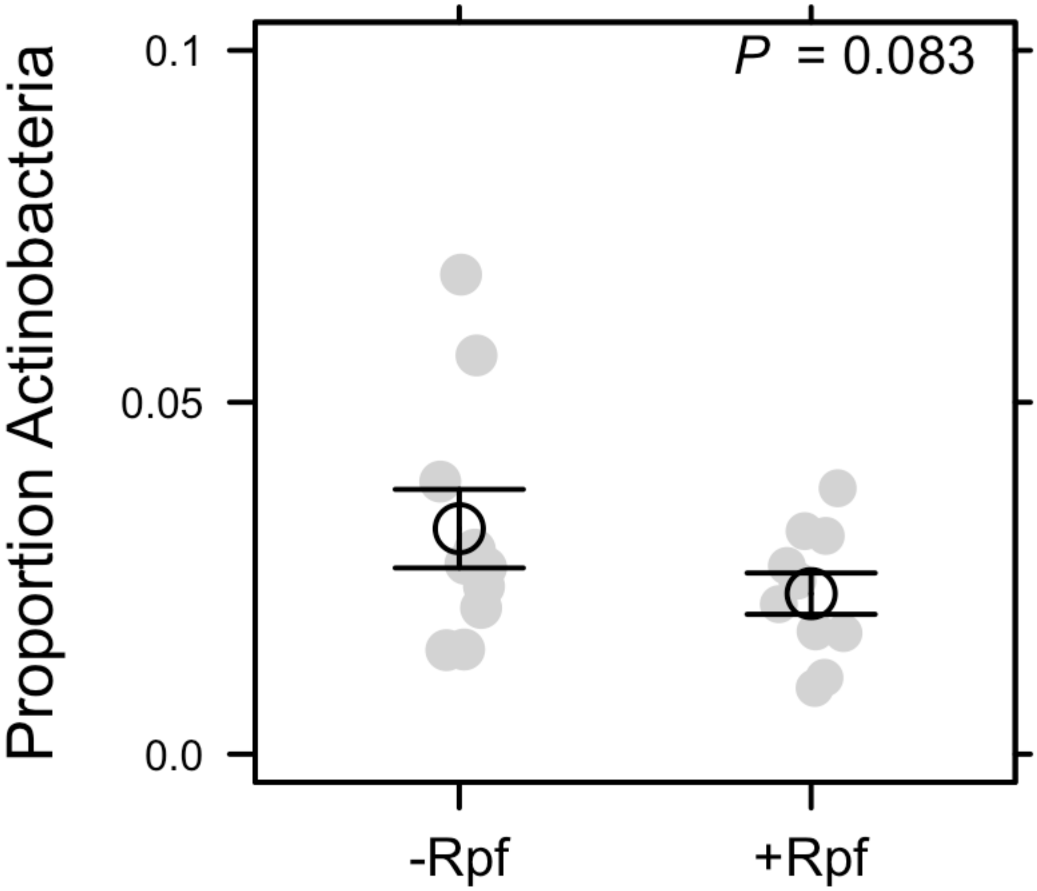

**Fig. S5**. Principal Coordinates Analysis (PCoA) demonstrating the effect of Rpf treatment on the active (RNA) and total (DNA) composition of Actinobacteria.

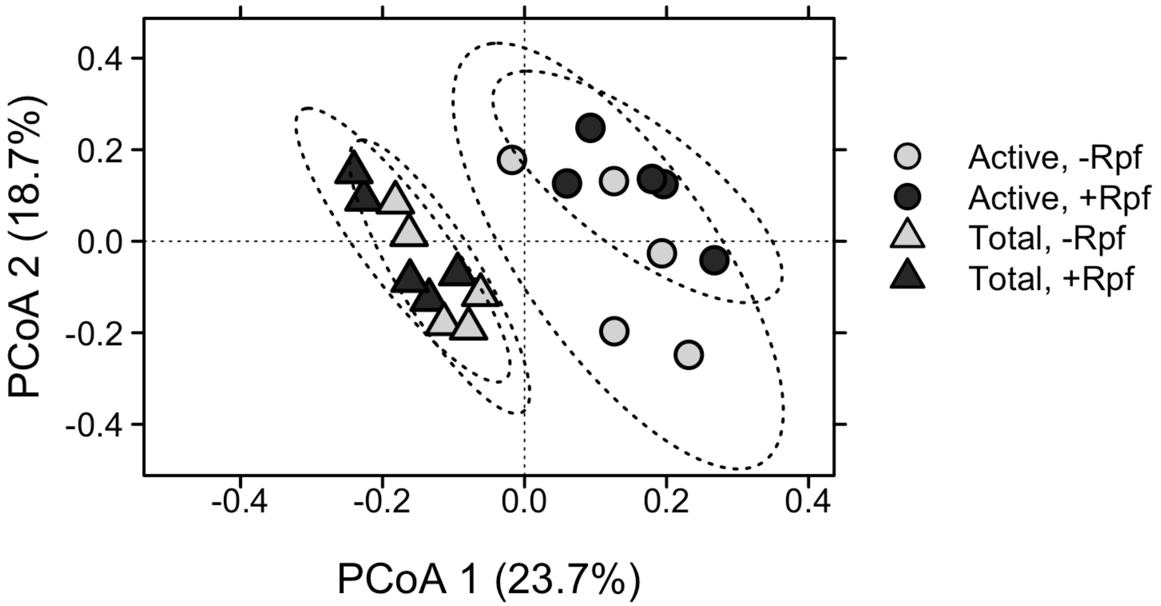

**Fig. S7**. Relative abundance of an unclassified OTU belonging to the Solirubrobacterales (Actinobacteria) that was identified in our indicator species analysis as having an association with the +Rpf treatment of the RNA pool. Data were relativized to the total number of actinobacterial sequences in a sample. Black symbols represent mean ± 95 % confidence intervals. Grey symbols represent individual observations.

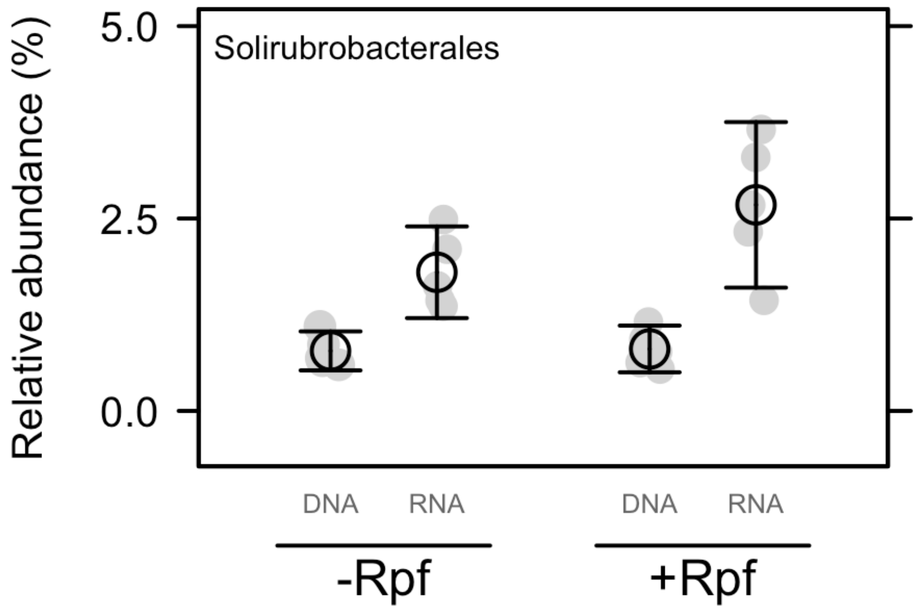

**Fig. S7**. To test for the effect of Rpf concentration on soil bacteria, we plated dilutions of soil onto Petri dishes containing R2A and measured the number of colony forming units (CFU) that formed following incubation. Briefly, we took soil cores (30 mm diameter x 115 mm depth) from Dunn Woods (Indiana), Griffy Woods (Indiana), and Machu Picchu forest (Peru). The soil samples were sieved through 2 mm sieve, homogenized by vortexing for 10 min, and partitioned into seven 0.5 g soil samples in 15 mL Falcon tubes. With the seven experimental units, we added enough recombinant Rpf to achieve the following concentration gradient: 0, 500, 1,000, 2,000, 4,000, 5,000, 8,700 nM. We then incubated the soils in the dark for three days at 25 °C. Following this incubation, we suspended each soil sample in 1 % pyrophosphate solution and vortexed vigorously for 30 m to separate bacterial cells from soil particles. We then plated 10-fold dilutions of the pyrophosphate solution in quadruplicate onto R2A plates containing cycloheximide (final concentration: 50 μg/mL) to inhibit fungal growth. The plates were incubated at 25 °C in the dark for one week and we counted CFUs using the Reichert Darkfield Quebec Colony Counter (ThermoFisher). Our results suggest that Rpf stimulates bacterial abundance at low to intermediate concentrations, but inhibits abundance at elevated concentrations. Error bars represent mean ± 1 $SE_{\bar{x}}$.

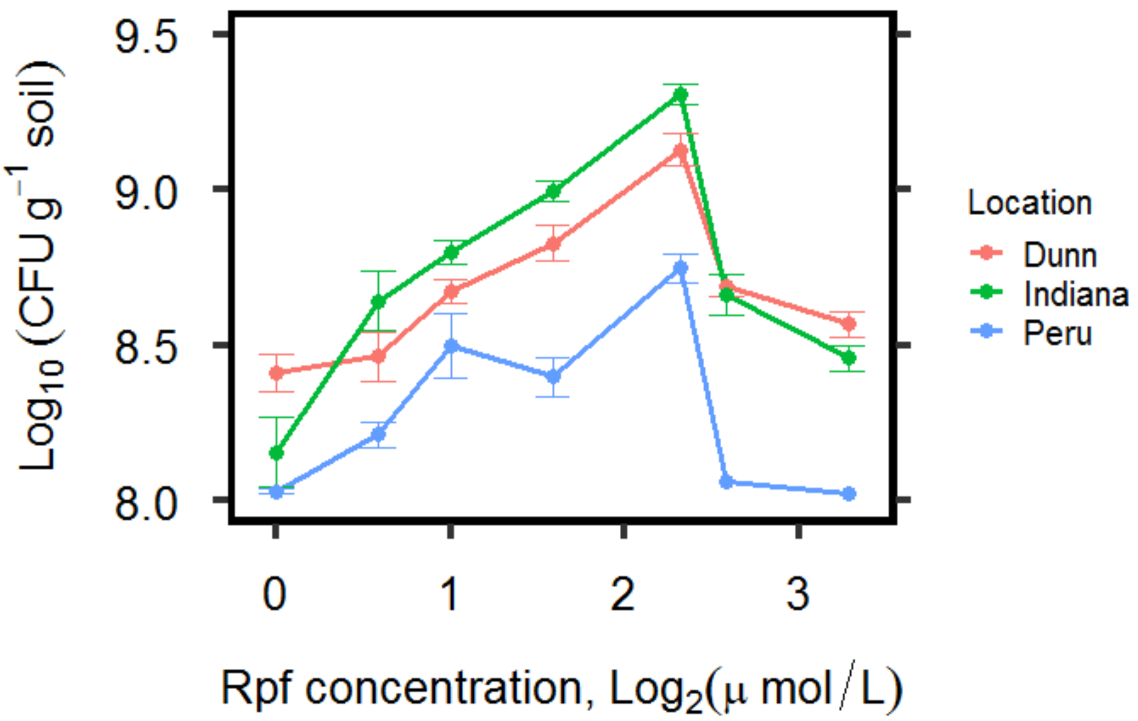

9.5
9.0
8.5
8.0
Log10 (CFU g-1 soil)
0
1
2
3
Rpf concentration, Log2(μ mol / L)
Location
Dunn
Indiana
Peru
